# Endogenous EWSR1-FLI1 degron alleles enable control of fusion oncoprotein expression in tumor cell lines and xenografts

**DOI:** 10.1101/2024.10.27.620498

**Authors:** James H. McGinnis, Alberto Bremauntz Enriquez, Florentina Vandiver, Xin Bai, Jiwoong Kim, Jessica Kilgore, Purbita Saha, Ryan O’Hara, Yang Xie, Laura A. Banaszynski, Noelle Williams, David G. McFadden

## Abstract

Pediatric malignancies frequently harbor chromosomal translocations that induce expression of fusion oncoproteins. The EWSR1-FLI1 fusion oncoprotein acts as a neomorphic transcription factor and is the dominant genetic driver of Ewing’s sarcoma. Interrogation of the mechanisms by which EWSR1-FLI1 drives tumorigenesis has been limited by a lack of model systems to precisely and selectively control its expression in patient-derived cell lines and xenografts. Here, we report the generation of a panel of patient-derived EWS cell lines in which inducible protein degrons were engineered into the endogenous EWSR1-FLI1 locus. These alleles enabled rapid and efficient depletion of EWSR1-FLI1. Complete suppression of EWSR1-FLI1 induced a reversible cell cycle arrest at the G_1_-S checkpoint, and we identified a core set of transcripts downstream of EWSR1-FLI1 across multiple cell lines and degron systems. Additionally, depletion of EWSR1-FLI1 potently suppressed tumor growth in xenograft models validating efforts to directly target EWSR1-FLI1 in Ewing’s sarcoma.

## INTRODUCTION

Fusion oncoproteins often act as singular drivers of tumor formation and maintenance in pediatric malignancies^1,2^. As such, these fusion proteins represent ideal targets for therapy, and exceptional responses have been observed across diverse malignancies driven by fusions encoding classically druggable proteins such as kinases^3–5^. However, many fusions encode proteins that act as neomorphic transcription factors, and therapeutic targeting of these proteins remains a challenge for drug discovery^6^. Ewing’s sarcoma (EWS), the second most common pediatric bone cancer, is characterized by the translocation of amino-terminal sequences of the RNA binding proteins EWSR1, FUS, or TAF15 with carboxy-terminal sequences that encode the DNA binding domain of an E-twenty six (ETS) transcription factor, most frequently FLI1^7^. The EWSR1-FLI1 fusion occurs in approximately 90% of EWS cases, and tumor genome sequencing efforts have identified few cooperating driver mutations^8–10^. These genetic studies suggest that EWSR1-FLI1 acts as the dominant, if not exclusive, driver of tumor initiation and growth in EWS. EWSR1-FLI1 therefore represents an ideal target for therapy. However, to date, viable small molecules have not been developed that directly or indirectly impair the function of EWSR1-FLI1^11^.

Since the discovery of EWSR1-FLI, investigators have sought to understand the mechanisms by which EWSR1-FLI1 initiates and drives tumorigenesis. Multiple independent studies have demonstrated that EWSR1-FLI1 binds to GGAA repeats within microsatellite sequences and induces transcriptional activation of neighboring genes^12–16^. Transcriptional activation by EWSR1-FLI1 is essential for the oncogenic function of the fusion^13^. Multiple studies have sought to understand the consequences of EWSR1-FLI1 suppression in EWS cell lines using RNA interference (RNAi) methods. These studies have reported varying phenotypes following suppression of EWSR1-FLI1 including no impact on proliferation, cell cycle arrest, senescence, or apoptosis even when using the same cell lines ^17–23^. For example, Smith et al. reported inhibition of soft agar colony formation, but no suppression of proliferation following knockdown of EWSR1-FLI1 in the A673 EWS cell line^19^. In contrast, Prieur et al. reported cell cycle arrest and apoptosis following EWSR1-FLI1 knockdown in A673 cells^17^. Which phenotypes represent on- or off-target effects of RNAi, or depend on the timing and/or completeness of EWSR1-FLI1 suppression remains unresolved.

A more recent study observed cell cycle arrest and apoptosis using CRISPR-Cas9 to delete regions including the junction of EWSR1 and FLI1. However, whether these phenotypes were induced by DNA damage or sgRNA off-target effects was not completely established^24^. Another study developed a system in which exogenous EWSR1-FLI1 fused to a degradation tag (dTag) was expressed in EWS cells in which endogenous EWSR1-FLI1 was subsequently genetically deleted^25^. While this system enabled specific depletion of EWSR1-FLI1 protein, phenotypes associated with EWSR1-FLI1 depletion were not reported.

Here we report the generation of a panel of patient-derived EWS cell lines in which we engineered orthogonal inducible protein degrons into the endogenous EWSR1-FLI1 locus. We employ these model systems to identify phenotypes following rapid and complete EWSR1-FLI1 depletion including cell cycle arrest, gene expression alterations, and potent suppression of EWS xenograft growth.

## RESULTS

### Degron tags enable depletion of endogenous EWSR1-FLI1

To identify phenotypes associated with EWSR1-FLI1 depletion in EWS, we developed an endogenous allele of EWSR1-FLI1 in the A673 cell line in which the auxin inducible degron (AID) was fused to the C-terminus of EWSR1-FLI1 using CRISPR/Cas9-mediated homologous recombination (Figure 1A, S1A). We successfully targeted AID into the EWSR1-FLI1 locus in the A673 cell line (referred to as A673 EF^AID^). The TIR1 E3 ligase from plants enables auxin (indoleacetic acid, IAA)-regulated proteasomal targeting of AID-tagged proteins^26^. We expressed *Oryza sativa* (rice) TIR1 in wild-type and EF^AID^-targeted A673 cells using a lentiviral vector. Treatment with IAA (100μM) induced complete (Figure 1B) and rapid (Figures 1C) degradation of endogenous EWSR1-FLI1-AID. Additionally, the EWSR1-FLI1 depletion was reversible, as removal of IAA from culture media resulted in recovery of EWSR1-FLI1 expression after 24 hours (Figure 1C). We also expressed modified TIR1 mutants (F74A and F74G) in A673 EF^AID^ cells to enhance the specificity of the AID system using potent chemically-modified auxin analogs^27^. Following expression of TIR1^F74A^ or TIR1^F74G^ in A673^EF-AID^ cells we observed complete depletion of EWSR1-FLI1 with 300nM 5-Ph-IAA (Figure S1B).

**Figure 1:**
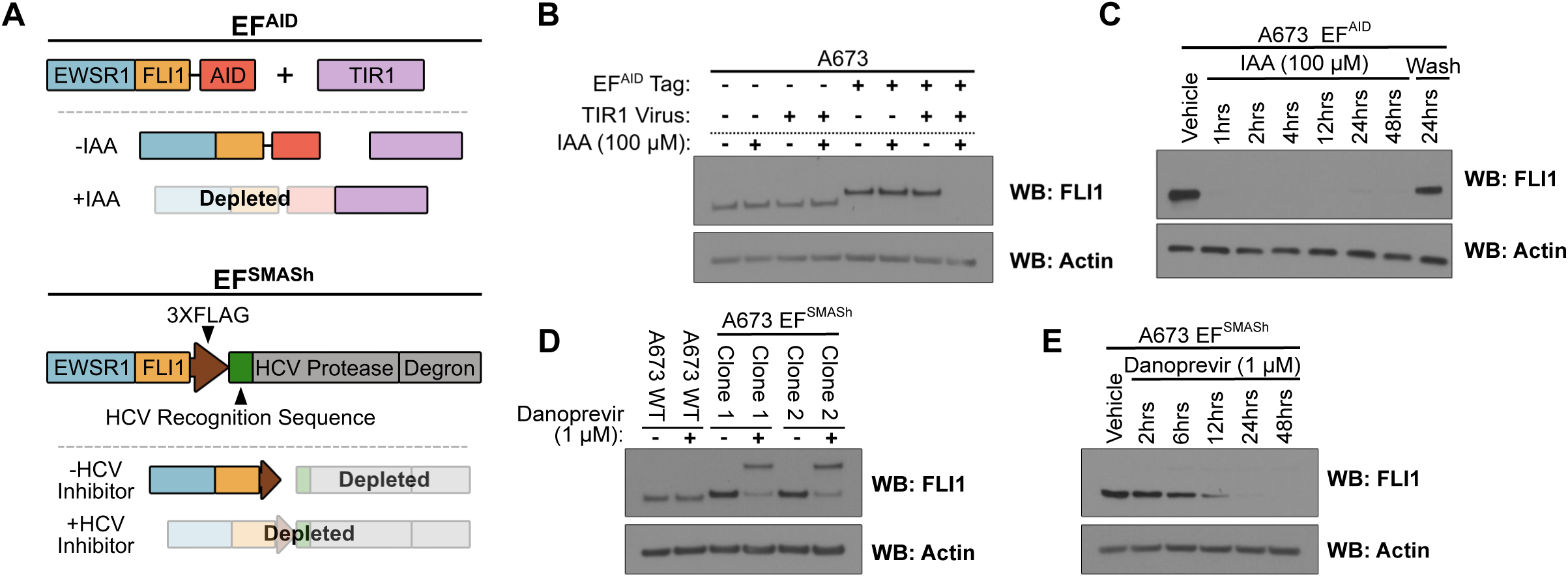
Degron tags enable depletion of endogenous EWSR1-FLI1. A) Schematic depicting SMASh and AID based degron approaches for depletion of endogenous EWSR1-FLI1. B) Immunoblot for EWSR1-FLI1 (FLI1) in indicated cell lines. Cell lines were exposed to either DMSO or IAA (100 μM) for 24 hours prior to collection. C) Immunoblot for EWSR1-FLI1 (FLI1) in indicated cell lines. Cell lines were exposed to either DMSO or IAA (100 μM) for the indicated time prior to collection. D) Immunoblot for EWSR1-FLI1 (FLI1) in indicated cell lines. Cell lines were exposed to either DMSO or danoprevir (1 μM) for 24 hours prior to collection. E) Immunoblot for EWSR1-FLI1 (FLI1) in indicated cell lines. Cell lines were exposed to either DMSO or danoprevir (1 μM) for the indicated time prior to collection.

We were not successful in targeting AID into the EWSR1-FLI1 locus in additional EWS cell lines including TC-32 and SK-N-MC. Although we obtained clones that harbored the EWSR1-FLI1-AID allele, in every case the wild-type, non-targeted, EWSR1-FLI1 locus was duplicated, suggesting strong selective pressure against the AID fusion in TC-32 and SK-N-MC cell lines (data not shown). Review of the literature suggested that A673 cells were less sensitive to RNAi-mediated suppression of EWSR1-FLI1 compared to other EWS cell lines^19^. We hypothesized that EWSR1-FLI1-AID fusion represented a hypomorphic variant of EWSR1-FLI1 that was tolerated only in A673 cells.

To develop degron EWSR1-FLI1 alleles in additional cell lines, we turned to the Small Molecule Assisted Shut-off (SMASh), a one-component system that was reported to enabled depletion of SMASh-tagged proteins^28^. The SMASh tag consists of a Hepatitis C viral (HCV) NS3 protease, NS3 cleavage recognition sequence, and degron (Figure 1A). Following translation of the SMASh-tagged protein, the HCV NS3 protease rapidly cleaves the NS3 recognition sequence which separates the protein of interest from the C-terminal NS3 protease and degron, leaving only a small six amino acid C-terminal peptide on the protein of interest. Upon treatment with an HCV protease inhibitor, cleavage of the HCV NS3 protease and degron is blocked. Therefore, all newly translated protein remains linked to the C-terminal degron and is therefore degraded. The kinetics of protein depletion represents a key difference between SMASh and AID. Upon treatment with auxin, all AID-tagged protein is rapidly degraded. In contrast, following treatment with NS3 inhibitors, the protein of interest that was already separated from the C-terminal degron is lost at its native half life.

We hypothesized that the self-cleaving nature of the SMASh system, which leaves a small C-terminal tag less likely to impair EWSR1-FLI1 function, would enable targeting of the degron into additional EWS cell lines. Indeed, we successfully targeted the SMASh degron into the endogenous EWSR1-FLI1 locus in A673, TC-32, and SK-N-MC cells (Figure S1C-D). We utilized multiple targeting vectors, some of which included a C-terminal epitope tag (Table 1). Interestingly, we obtained targeted TC-32 and SK-N-MC cells exclusively with constructs that omitted the epitope tags, similar to our experience targeting AID into these cell lines. These findings were consistent with studies suggesting the C-terminal sequences of EWSR1-FLI1 contributed to the oncogenic function of the fusion, and again raised the possibility that larger C-terminal fusions compromised EWSR1-FLI1 function^29,30^.

**Table 1:** Endogenous degron cell lines.

| **Table 1: Endogenous degron cell lines** |  |  |
| --- | --- | --- |
| Cell Line | Degron | Epitope tag |
| A673 | SMASh | 3XFLAG |
| A673 | SMASh | V5 |
| A673 | SMASh | None |
| A673 | AID | 3XFLAG |
| A673 | AID | None |
| TC-32 | SMASh | None |
| SK-N-MC | SMASh | None |

We treated EF^SMASh^ cell lines with the HCV NS3 protease inhibitor danoprevir (1μM) and observed near-complete depletion of EWSR1-FLI1 at 24hrs, consistent with the reported half life of the EWSR1-FLI1 fusion protein (Figure 1D, S1D)^31^. We variably observed (compare Figure 1D and 1E) accumulation of a higher molecular weight FLI1 band following danoprevir treatment that was consistent with incomplete degradation of the retained SMASh tag on the EWSR1-FLI1 protein (EWSR1-FLI1-SMASh). We confirmed the identity of the high molecular weight band by blotting for the Myc epitope tag engineered downstream of the NS3 cleavage site (Figure S1E). We did not detect a 30kDa band, the expected molecular weight of the cleaved HCV NS3 protease-degron tag, suggesting that depletion of the cleaved SMASh tag was more efficient than EWSR1-FLI1-SMASh. We concluded that EWSR1-FLI1-AID and EWSR1-FLI1-SMASh alleles enabled control of endogenous EWSR1-FLI1 levels in patient-derived EWS cell lines, albeit with different depletion kinetics.

### C-terminal AID tag on EWSR1-FLI1 does not disrupt DNA binding

Several groups have shown that EWSR1-FLI1 recognizes and binds GGAA motifs and GGAA repeats within microsatellite regions through its ETS DNA binding domain encoded in the C-terminal fusion partner FLI1 ^14–16^. We performed CUT&RUN analysis of EWSR1-FLI1 binding in wild-type A673 and A673 EF^AID^; TIR1 cells to determine if the C-terminal AID tag impacted DNA binding. Analysis of EWSR1-FLI1 peaks demonstrated that EWSR1-FLI1 and EWSR1-FLI1-AID bound highly overlapping regions (Figure 2A). Ranking the EWSR1-FLI1 peaks by intensity in the A673 parental line, we observed strong concurrence between FLI1 peaks in A673 and A673 EF^AID^; TIR1^F74A^ cell line (Figure 2B).

**Figure 2:**
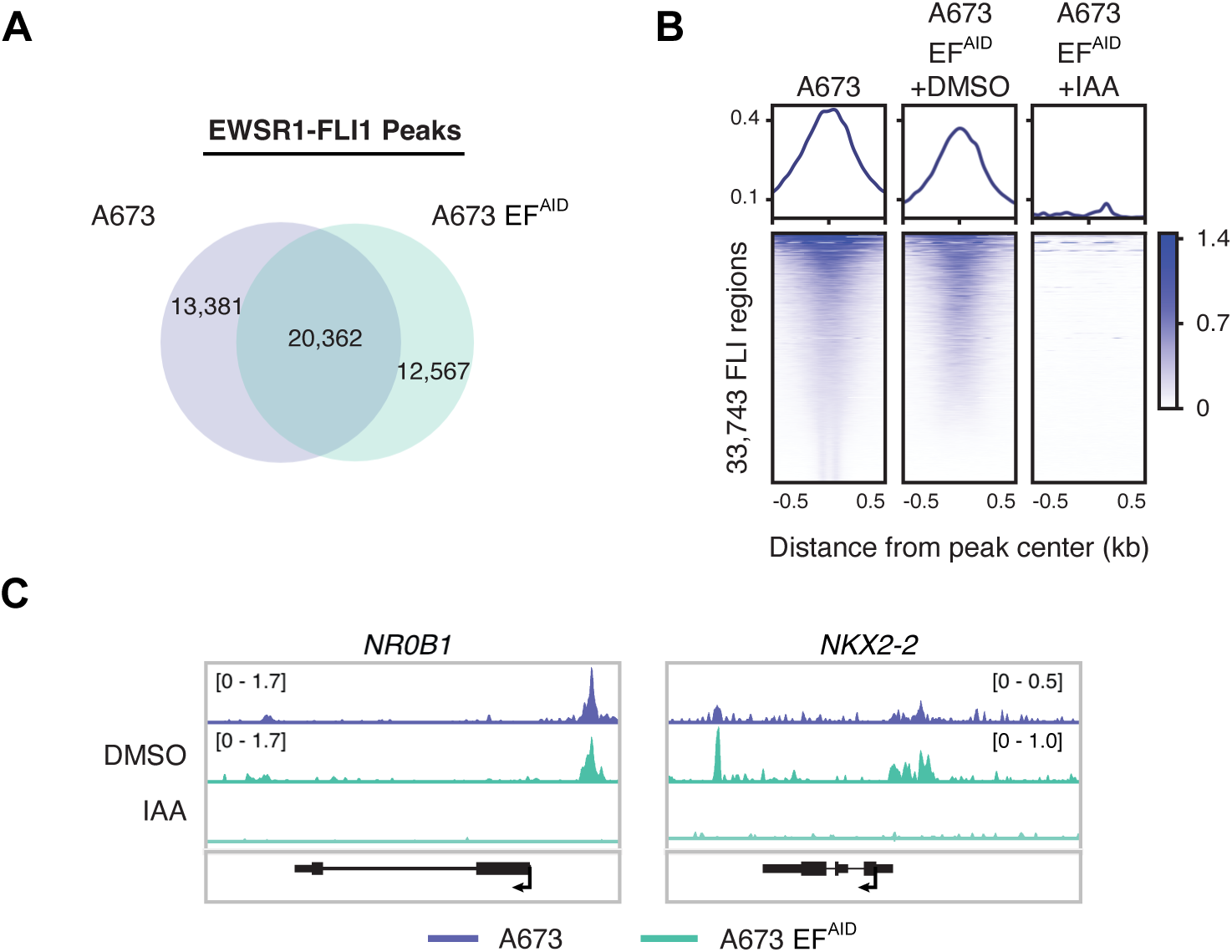
C-terminal AID tag on EWSR1-FLI1 does not disrupt DNA binding. A) Venn diagram representing overlap of FLI enriched regions identified from A673 and A673 EF^AID^;TIR1 F74A cells B) Average profiles (top) and heatmaps (bottom) of FLI CUT&RUN enrichment at FLI enriched regions in A673 cells (n=33,743) in A673 cells and A673 EF^AID^;TIR1 F74A cells treated with DMSO or 5-Ph-IAA (300 nM) for 24 hrs. 0.5 kb around the peak center are displayed for each analysis C) Genome browser representations of FLI CUT&RUN in A673 cells and A673 EF^AID^;TIR1 F74A cells treated with either DMSO or 5-Ph-IAA (300 nM) for 24 hrs. The y-axis represents read density in reads per million mapped reads (rpm).

We also performed binding site motif analysis of EWSR1-FLI1-bound peaks. We observed similar enrichment of single GGAA (FLI1) and GGAA multimers within microsatellites (EWS:FLI) motifs in A673 and A673 EF^AID^ cells (Figure S2A). These datasets suggested that addition of the AID tag did not impair EWSR1-FLI1 binding to DNA and suggested that the potential hypomorphic nature of C-terminal AID fusions were likely independent of direct DNA binding, consistent with prior studies reporting a DNA-binding-independent function of the EWSR1-FLI1 C-terminal sequences^29,30^.

We also utilized CUT&RUN to determine if 5-Ph-IAA induced efficient depletion of chromatin bound EWSR1-FLI1 in A673 EF^AID^;TIR1^F74A^ cells. Following treatment with 300nM 5-Ph-IAA for 24 hours, we observed complete loss of EWSR1-FLI1 peaks (Figure 2B). We manually reviewed datasets for several direct EWSR1-FLI1 target genes including *NR0B1, NKX2-2, CCND1,* and *VRK1*. We observed strong EWSR1-FLI1 peaks at these loci in parental A673 and A673 EF^AID^;TIR1^F74A^ cell lines (Figure 2C, S2B). Consistent with the global analysis of EWSR1-FLI1 peaks, and western blotting from whole cell lysates, complete loss of EWSR1-FLI1 peaks was observed following 5-Ph-IAA treatment. We concluded that AID enabled efficient degradation of nuclear chromatin-bound EWSR1-FLI1-AID.

### EWSR1-FLI1 suppression induces G_1_-S cell cycle arrest

We next sought to establish reliable phenotypes following acute loss of EWSR1-FLI1 in EWS cells. We treated EF^SMASh^ cell lines and wild-type parental cell lines with DMSO or danoprevir (1μM) for 6 days and monitored proliferation by cell counting. Parental cell lines exhibited no change in proliferation in response to danoprevir. In contrast, danoprevir reduced proliferation in EF^SMASh^ cell lines (Figure 3A and S3A). We also observed reduced proliferation following depletion of EWSR1-FLI1 in A673 EF^AID^ cells following treatment with IAA. In contrast, we did not observe a change in proliferation in wild-type A673 expressing TIR1 (to assess for AID-independent, auxin-dependent TIR1 effects), or in A673 EF^AID^ without TIR1 expression following IAA treatment (Figure 3B, S3B). We noted a greater suppression of proliferation in TC32 EF^SMASh^ and SKNMC EF^SMASh^ cell lines following danoprevir treatment compared to A673 EF^SMASh^ and A673 EF^AID^ which we suspected was consistent with less EWSR1-FLI1 dependence of A673 cells as discussed above.

**Figure 3:**
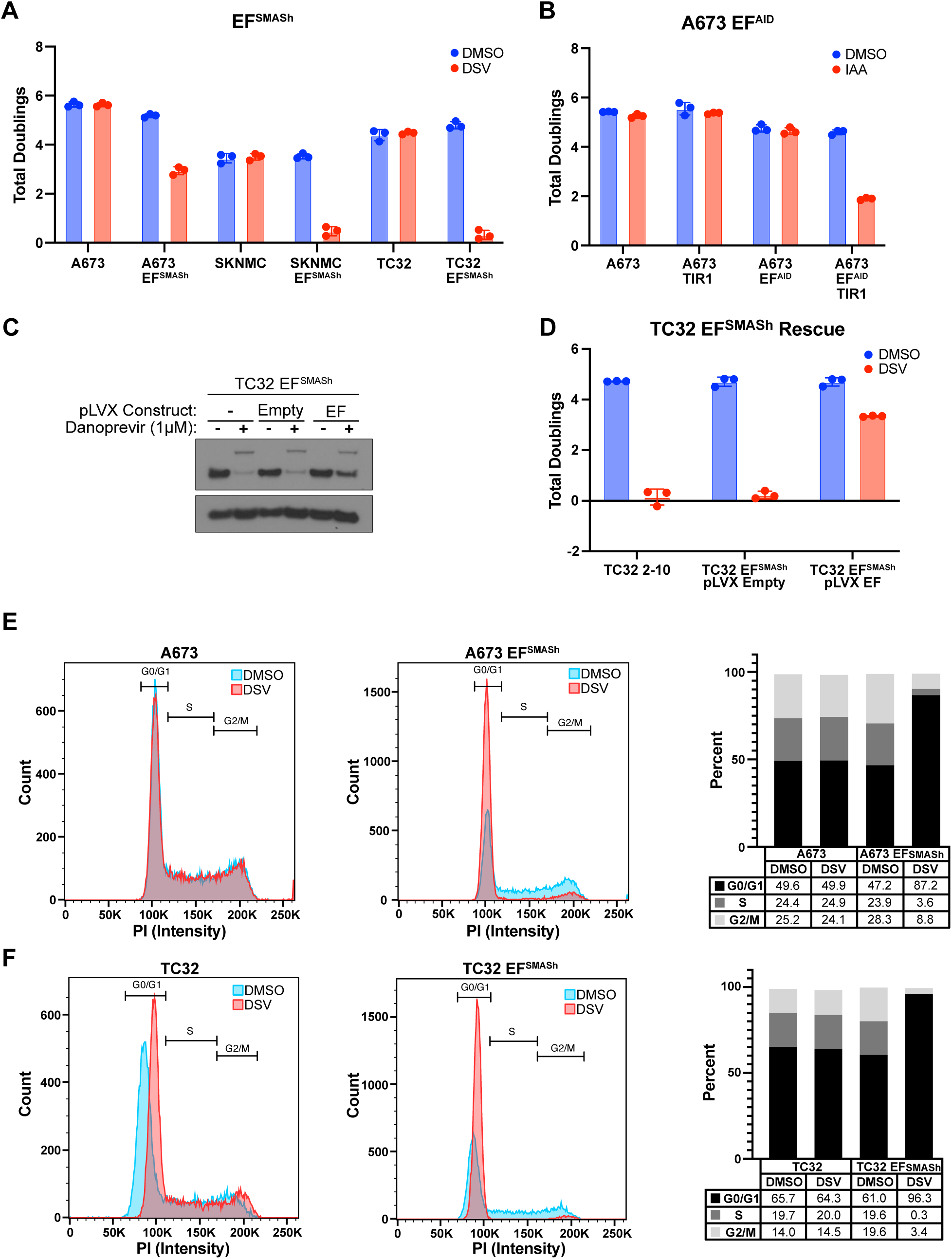
EWSR1-FLI1 depletion induces G1/S arrest. A) Population doublings after 6 days of treatment. Indicated cell lines were exposed to vehicle or 1 μM danoprevir (DSV). B) Population doublings after 6 days of treatment. Indicated cell lines were exposed to DMSO or IAA (100 μM). C) Immunoblot for EWSR1-FLI1 (FLI1) in TC32 EF^SMASh^ cells with indicated pLVX constructs. Cell lines were exposed to either Vehicle or danoprevir (1 μM) for 24 hours prior to collection. D) Population doublings after 6 days of treatment. TC32 EF^SMASh^ cells with indicated pLVX constructs (C) were exposed to vehicle or 1μM danoprevir. E-F) Cell cycle analysis using propidium iodide. Flow cytometry plots (left two panels) for indicated cell lines treated with either vehicle or 1 μM danoprevir for 72 hours before cells were collected. Plot (left panel) of cells (percentage) in each phase of the cell cycle based on flow cytometry data.

The variable retention of the EWSR1-FLI1-SMASh-degron high molecular weight protein following danoprevir treatment in EF-SMASh cell lines (See Figures 1C, S1C, 3C) raised the possibility that the suppression of cellular proliferation observed following EWSR1-FLI1 depletion could be due to dominant-negative action of the retained EWSR1-FLI1-SMASh-degron. We therefore generated TC32 EF^SMASh^ cells that constitutively and exogenously expressed wild-type EWSR1-FLI1. We observed near physiological levels of EWSR1-FLI1 in these cells (Figure 3C). We treated cells with danoprevir (1μM) to induce depletion of endogenous EWSR1-FLI1-SMASh and followed cell proliferation for multiple cell doublings. We observed rescued proliferation following depletion of endogenous EWSR1-FLI1 with danoprevir (Figure 3D, S3C). We concluded that loss of endogenous EWSR1-FLI1 was responsible for impaired proliferation in EF^SMASh^ cell lines following danoprevir treatment, as opposed to a dominant-negative activity of retained high molecular weight EWSR1-FLI1-SMASh protein.

To characterize impaired proliferation following EWSR1-FLI1 depletion, we performed cell cycle analysis with propidium iodide. A673^EF-SMASh^, TC-32^EF-SMASh^, and SKNMC^EF-SMASh^ and parental cell lines were treated with 1μM danoprevir treatment for 48 hours. We observed an accumulation of cells in the G_1_-S phase following danoprevir treatment that was not observed in non-targeted parental cell lines cells (Figure 3E-F and Figure S3D). The reproducibility of this phenotype across multiple cell lines suggested that cell cycle arrest at the G_1_-S checkpoint was responsible for reduced proliferation of EWS cells following EWSR1-FLI1 depletion.

One study observed increased markers of cellular senescence following EWSR1-FLI1 suppression with siRNA^18^. We leveraged the reversibility of SMASh and AID EWSR1-FLI1 degron systems following washout of danoprevir or IAA, respectively, to determine whether cell cycle arrest following EWSR1-FLI1 depletion was reversible. We observed recovery of proliferation within 24 hours following the removal of danoprevir or IAA from the growth media of A673 EF^SMASh^ and A673 EF^AID^;TIR1 cell lines (Figure S2E-F). We concluded from these studies that EWSR1-FLI1 depletion induced a reversible cell cycle arrest as opposed to irreversible cellular senescence.

### Transcriptome profiling following EWSR1-FLI1 suppression identifies a core set of EWSR1-FLI1 regulated genes

We next assessed the specificity and reproducibility of transcriptional programs following EWSR1-FLI1 depletion across independent cell lines and degrons. We performed transcriptome sequencing 24 hours following depletion of EWSR1-FLI1 with danoprevir (1μM) in two independently generated A673 EF^SMASh^, TC-32 EF^SMASh^, and SKNMC EF^SMASh^ cell lines (Figure 4A, Table S1). These studies revealed transcriptional heterogeneity following EWSR1-FLI1 depletion in different cell lines (Figure 4B). We compared transcriptional responses following EWSR1-FLI1 depletion in each of the SMASh cell lines to reported gene sets using Gene Set Enrichment analysis (GSEA)^3233^. We consistently identified gene sets associated with suppression of EWSR1-FLI1 in EWS cell lines, or those in which EWSR1-FLI1 was exogenously expressed in mesenchymal progenitor cells (Table S2-S3).

**Figure 4:**
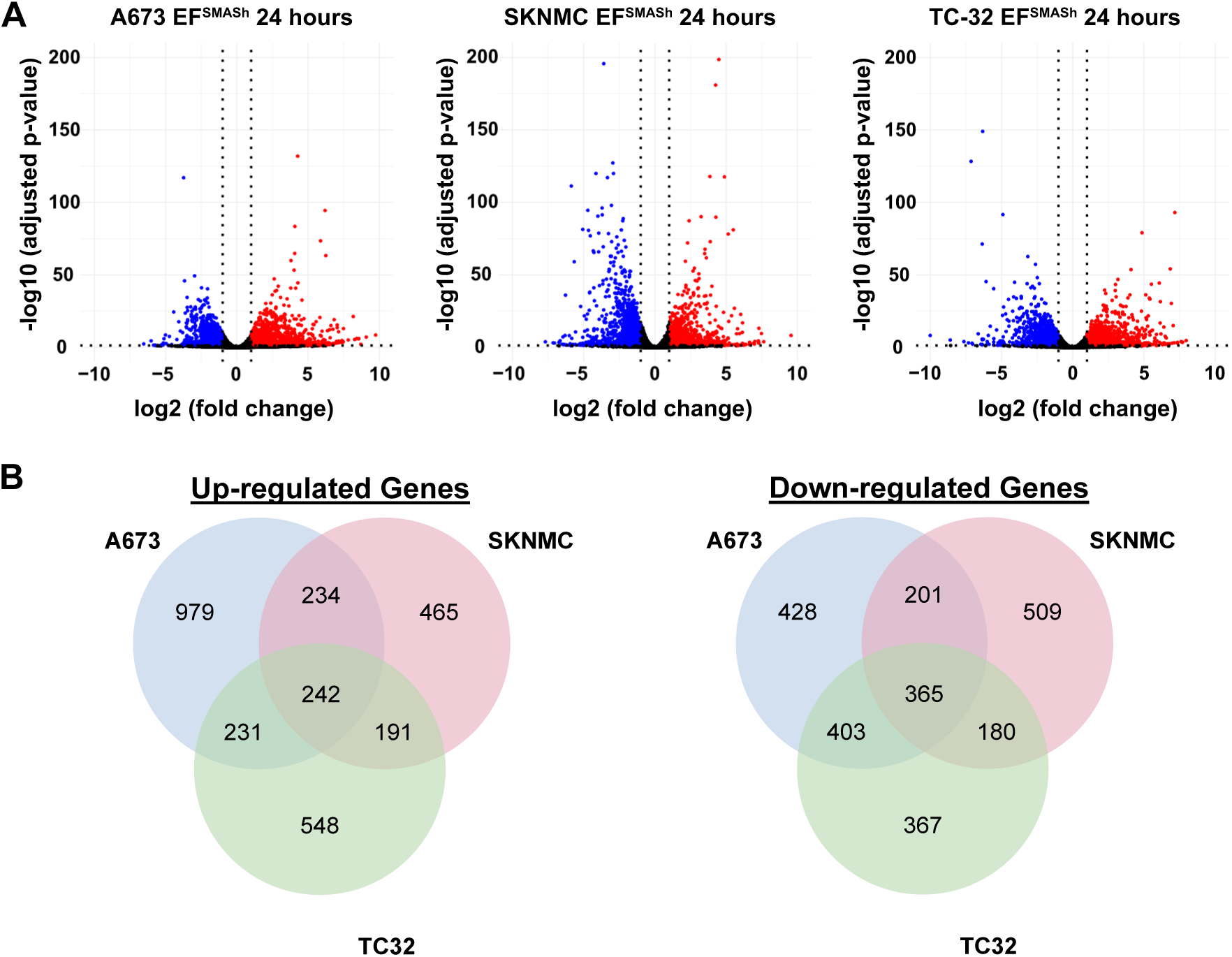
Core set of EWSR1-FLI1 response genes shared across EWS cell lines. A) Volcano plots of RNA sequencing data obtained from indicated cell lines following treatment with 1 μM danoprevir. Red dots represents genes that are significantly differentially induced. Blue dots represents genes that are significantly differentially repressed. B) Venn diagram comparing up-regulated genes (left) or down-regulated genes (right) across the three cell lines tested.

We hypothesized that transcriptional targets essential for the oncogenic function of EWSR1-FLI1 would be conserved across cell lines. We therefore defined an ‘EF core signature’ of up- (n = 242) and down- (n = 365) regulated transcripts following EWSR1-FLI1 depletion in A673, SK-N-MC, and TC-32 cells engineered with the SMASh degron (Figure 4B, Table S4). We performed overrepresentation analysis using the EF core signature transcripts to identify pathways and processes associated with suppression of EWSR1-FLI1 function. We again observed gene sets associated with modulation of EWSR1-FLI1 in EWS and mesenchymal progenitor cells, as well as several signatures associated with cancer cell proliferation, stemness, and invasiveness (Table S5-S6).

We also performed transcriptome sequencing following EWSR1-FLI1 depletion in A673 EF^AID^ and A673 EF^SMASh^ cell lines to compare the SMASh and AID degron systems (Figure S4A). Comparing transcripts with a 2-fold increase or decrease following degron induction, we observed a strong correlation between the two gene sets (R = 0.838, p = 3.29 × 10^-161^, Figure S4B). These data suggested the dominant transcriptional alterations following depletion of EWSR1-FLI1 were driven by loss of the fusion protein as opposed to nonspecific alterations due to the individual degron systems or the chemical inducers of depletion (danoprevir or IAA).

Finally, we profiled transcriptome dynamics following EWSR1-FLI1 suppression at 6, 12, and 24 hours following administration of danoprevir (312nM) in A673 EF^SMASh^ and A673 WT cells (Table S7). We noted an approximately 6-hour half-life of EWSR1-FLI1 protein following treatment with danoprevir in A673 EF^SMASh^ cells (Figure 1E). We observed alterations in the EF core signature transcripts at 6hrs following danoprevir administration, suggesting a ∼50% reduction in EWSR1-FLI1 protein was sufficient to impair EWSR1-FLI1 transcriptional function (Figure S4C, D). This finding is consistent with other studies demonstrating that a narrow window of EWSR1-FLI1 expression level is required for oncogenic function^34,35^. We noted increased amplitude of EF core signature transcript levels at 12- and 24-hour timepoints (Figure S2C-D). Finally, analysis of RNA sequencing datasets of A673 WT cells treated with danoprevir demonstrated few nonspecific transcriptional changes highlighting the exceptional specificity of this degron system (Figure S4E-F).

### EWSR1-FLI1 is required for tumor growth in a xenograft model

We evaluated whether SMASh- and AID-mediated EWSR1-FLI1 depletion was feasible in animal models using cell line xenografts. Danoprevir was developed for treatment of hepatitis C, including optimization for concentration in liver^36^. We were concerned the optimization of danoprevir for liver concentration would impair drug delivery to tumors. We therefore examined danoprevir levels in immunodeficient mice bearing A673 EF^SMASh^ xenograft. After xenograft tumors reached >200 mm^3^ (range 234-450 mm^3^) danoprevir was administered by oral gavage at 30 mg/kg twice daily for 2 days. Mice were sacrificed 3 hours after the final dose and danoprevir levels in plasma, tumor and liver were determined by LC-MS/MS. Consistent with prior studies, we observed accumulation of danoprevir in the liver and very low concentration in xenografts, suggesting the drug exposure in tumor models was below the concentration required for efficient SMASh-mediated EWSR1-FLI1 depletion (Figure S5A). Indeed, we did not observe EWSR1-FLI1 depletion in tumor protein lysates following danoprevir treatment (data not shown).

We next evaluated AID-mediated EWSR1-FLI1 depletion in animal models. We established A673 EF^AID^ xenografts in immunodeficient mice, and initiated IAA treatment after tumors reached 400mm^3.^ IAA was administered at 200 mg/kg twice daily IP and animals were sacrificed after 9 doses. We observed tumor regression in mice treated with IAA at 2 days or 4 doses (35% decrease in tumor volume) and sustained regression to the study endpoint of 5 days or 9 doses (57% decrease in tumor volume) (Figure S5B). IAA treatment was well tolerated, as we observed no change in mouse weight over the 9 doses (Figure S5C). To determine IAA tumor exposure mice were sacrificed at 6, 12, and 24 hours after the final dose of IAA. IAA concentrations were determined in the plasma and tumor by LC-MS/MS. We observed good exposure of IAA in the tumor and plasma at all time points tested (Figure S5D). Concentrations of IAA in tumors were above 558.8 ng/g (3.25 µM) 24 hours following the final dose.

We treated an independent cohort of A673 EF^AID^ xenograft-bearing animals with IAA to evaluate EWSR1-FLI1 protein depletion and extend analysis of the tumor response. We also implanted A673 EF^AID^ cells that lacked TIR1 expression to confirm IAA treatment did not impact tumor growth in an AID-independent manner. Tumor bearing mice were treated with vehicle or IAA 200 mg/kg twice daily until vehicle treated mice met the criteria for euthanasia, a total of 11 days. A673 EF^AID^ xenografts treated with IAA exhibited an initial decrease in tumor volume, similar to that observed in the pilot study. However, xenografts subsequently increased in size and grew at a similar rate to vehicle treated tumors (Figure 5A).

**Figure 5:**
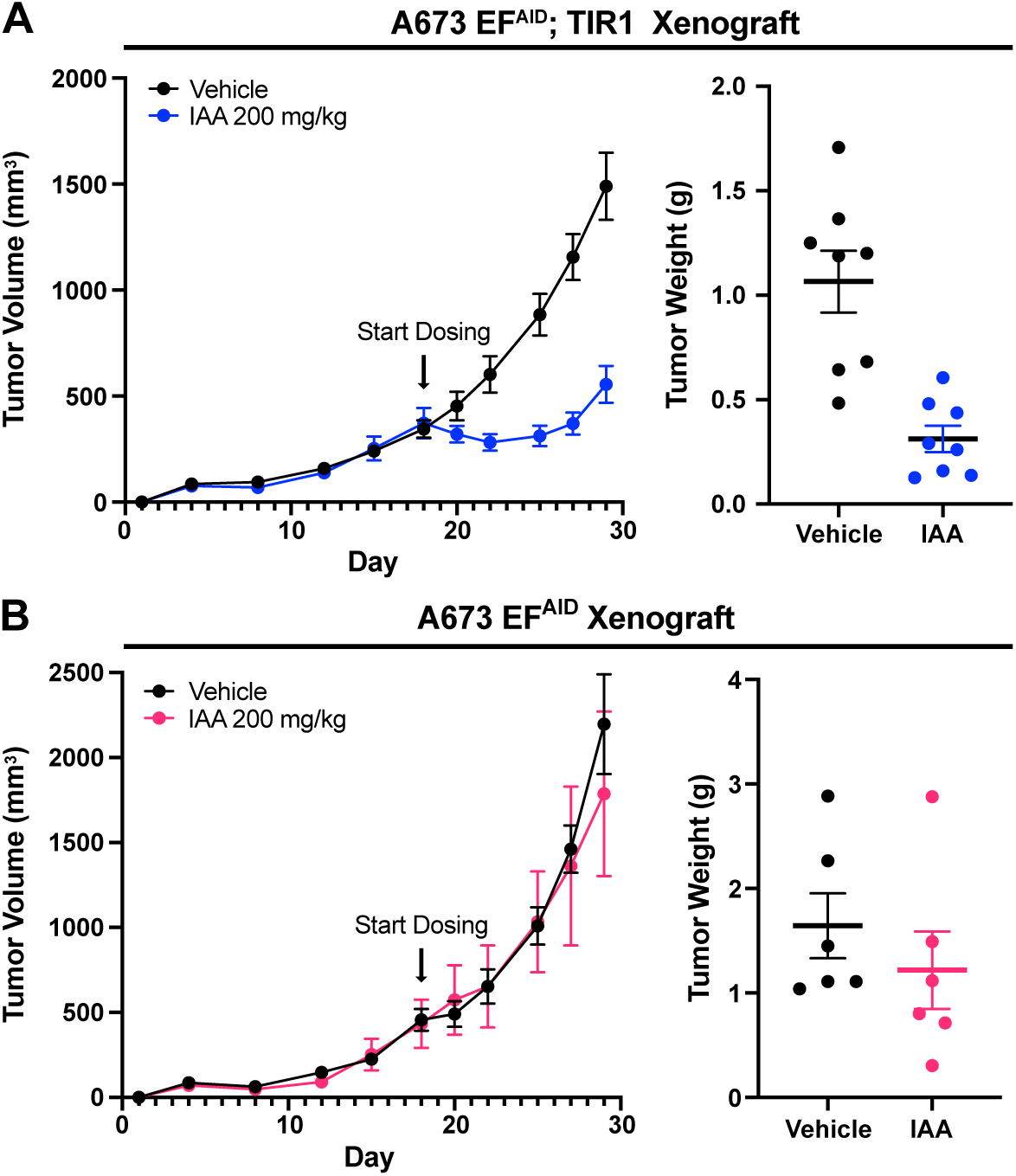
EWSR1-FLI1 is required for tumor maintenance in vivo. A) Tumor volume (left) and tumor mass (right) for A673 EF^AID^;TIR1 xenografts treated with vehicle (n = 8) or 200 mg/kg IAA (n = 8). Vehicle or IAA was administered twice daily via oral gavage for a total of 11 days. B) Tumor volume (left) and tumor mass (right) for A673 EF^AID^: xenografts treated with vehicle (n = 6) or 200 mg/kg IAA (n = 6). Vehicle or IAA was administered twice daily via oral gavage for a total of 11 days.

We noted no difference in the growth of IAA-treated A673 EF^AID^ xenografts that did not express TIR1 compared to vehicle control, suggesting that IAA did not impact tumor growth through an AID-independent mechanism (Figure 5B). Mice tolerated 11 days of IAA treatment without evidence of weight loss (Figure S5E-F).

We examined EWSR1-FLI1 protein levels in xenograft lysates collected from mice sacrificed at D2, D4 and D11. A673 EF^AID;TIR1^ tumor lysates obtained from vehicle or IAA treated mice on D2 showed depletion of EWSR1-FLI1 with IAA treatment (Figure S5G). However, tumors collected on D4 showed appreciably less depletion of EWSR1-FLI1 following IAA treatment. Comparing EWSR1-FLI1 levels between DMSO and IAA treatment at D11, we observed no depletion of EWSR1-FLI1 in IAA-treated samples. This result raised the possibility of technical escape from AID-mediated depletion. We considered loss of TIR1 expression in A673 EF^AID^ cells as one potential mechanism of technical escape. A673 EF^AID^ xenografts were generated from one A673 EF^AID^ clone that was subsequently transduced with TIR1-expressing virus. We did not regenerate single cell clones following transduction with the TIR1-expressing virus, raising the possibility of selection for cells expressing lower levels of TIR1 that would be expected to retain EWSR1-FLI1-AID expression in the presence of IAA. We observed similar TIR1 protein expression in DMSO and IAA samples at D2 and D4 (Figure S5H). However, xenografts harvested after 11 days of dosing exhibited decreased TIR1 expression compared to controls. We concluded that EWSR1-FLI1 depletion suppressed xenograft growth and that a strong selective pressure against EWSR1-FLI1 loss in xenografts promoted emergence of A673 EF^AID^ cells with low TIR1 expression that retained EWSR1-FLI1 expression.

## DISCUSSION

Here we report two orthogonal systems that enabled control of endogenous EWSR1-FLI1 expression in patient-derived EWS models. These endogenous alleles engineered that harbored C-terminal inducible protein degrons enabled rapid and specific depletion of EWSR1-FLI1 in cell lines and subcutaneous xenografts.

Our investigation of EWSR1-FLI1 function differed in strategy from previous studies that have relied on RNA interference techniques to deplete EWSR1-FLI1. We consistently observed cell cycle arrest at the G_1_-S checkpoint, consistent with some, but not all prior studies. The reversible nature of these degrons further enabled us to determine that cell cycle arrest following EWSR1-FLI1 depletion was reversible following restoration of fusion protein expression. Although this represents an artificial system, re-activation of the cell cycle argues against irreversible cellular senescence or apoptosis following EWSR1-FLI1 suppression.

Nonetheless, limitations exist with endogenous oncoprotein degron systems. First, for our EF^SMASh^ alleles variably retain EWSR1-FLI1-SMASh protein (Figure 1D) following danoprevir treatment. Retention of the EWSR1-FLI1-SMASh protein raised the possibility that this protein could a) retain hypomorphic function, or b) exert dominant-negative effects in EWS cells. However, the observation of G_1_-S checkpoint arrest and overlapping transcriptional changes using the orthologous EF^AID^ allele suggested that the phenotypes observed following SMASh-mediated depletion were due to loss of EWSR1-FLI1, rather than retention of EWSR1-FLI1-SMASh. Furthermore, rescue of proliferation following danoprevir-induced EWSR1-FLI1 depletion in TC32 EF^SMASh^ cells that exogenously express EWSR1-FLI1 strongly argues against a dominant-negative action of the retained EWSR1-FLI1-SMASh protein.

Throughout our experiments, A673 cells exhibited a higher tolerance for manipulation of endogenous EWSR1-FLI1. We successfully engineered A673 cells with larger C-terminal fusions, including AID, and SMASh fusions that encoded epitope tags. In addition, although we observed G_1_-S arrest in A673 EF^SMASh^ and A673 EF^AID^ cells following EWSR1-FLI1 depletion, arrest was not as complete as that observed in TC-32 EF^SMASh^ or SK-N-MC EF^SMASh^ cells. These findings are consistent with Smith et al., in which suppression of EWSR1-FLI1 in A673 cells did not impact proliferation in standard tissue culture^19^. However, in contrast to Smith et al., we observed a strong decrease in proliferation and induction of cell cycle arrest. We hypothesize that the complete and rapid loss of EWSR1-FLI1 in our models might account for this difference in phenotype following EWSR1-FLI1 suppression. However, other studies have also reported decreased proliferation following RNAi-mediated suppression of EWSR1-FLI1 in A673 cells^17^. Therefore, the different responses to EWSR1-FLI1 suppression in A673 cells remain ambiguous. Nonetheless cumulatively these data suggest that A673 cells might be less reliant on EWSR1-FLI1 to drive proliferation. A673 cells are known to harbor a BRAF^V600E^ mutation which is not observed in patient tumors^10^. Whether this mutation contributes to EWSR1-FLI1 independence in A673 cells is unknown.

We noted very few nonspecific transcriptional changes in wild-type EWS cells treated with the NS3 protease inhibitor (danoprevir) used to induce SMASh-mediated protein depletion (Figure S4). This specificity likely reflects the extensive drug development efforts invested in these compounds. Such specificity minimized potential off-target changes in gene expression that are frequently observed when using RNAi methods. Using multiple SMASh-targeted cell lines, we defined a conserved core gene expression signature following EWSR1-FLI1 depletion. We propose this core signature as a potential template for identification of the key effectors of the EWSR1-FLI1 oncogenic program.

We successfully employed inducible protein degradation in xenograft models using the EF^AID^ allele in A673 cells. We unexpectedly observed tumor regression following suppression of EWSR1-FLI1 in A673 EF^AID^ xenografts. Although A673-EF^AID^ xenografts rapidly became resistant to IAA, we observed technical escape from AID-mediated EF depletion driven by loss of TIR1 expression. The emergence of cells lacking TIR1 expression is consistent with a strong selective pressure against loss of EWSR1-FLI1 expression during EWS tumor growth. This result establishes a requirement for EWSR1-FLI1 expression in tumor maintenance. The observation of tumor regression also raises the possibility that the response to suppression of EWSR1-FLI1 differs in vivo compared to cell culture, even in immunocompromised hosts. Additional studies using these models will be required to define the mechanisms underlying the transient regression of xenograft tumors following EWSR1-FLI1 suppression.

These models might be especially useful to investigate the sensitivity of EWS cells to partial suppression of EWSR1-FLI1 function, which some studies have suggested enhances cellular migration and metastatic potential^37^. The specific domains and regions of EWSR1-FLI1 that are necessary and sufficient to drive and maintain tumor formation remain incompletely characterized^12,30,38,39^. We propose these endogenous degron allele models as one potential avenue to perform structure-function studies of EWSR1-FLI1. The simplicity of exogenous expression of various truncations and mutants followed by depletion of endogenous EWSR1-FLI1 represents a straightforward assay of EWSR1-FLI1 function that could be employed in cell culture or xenograft assays (see Figure 3D). The identification of shared phenotypes for EWSR1-FLI1 depletion across cell lines, namely G_1_-S arrest and a core set of altered transcripts, provide a clear functional readout for future investigation and potential screening readouts. We propose this collection of EWS cell lines with tunable endogenous EWSR1-FLI1 degron alleles as an important extension of the available model systems to facilitate detailed interrogation of fusion oncoprotein function and accurate modeling of acute therapeutic inhibition of these powerful oncoproteins, exemplified by EWSR1-FLI1.

## RESOURCE AVAILABILITY

### Material availability

All plasmids reported in this manuscript are available upon request or through Addgene.org.

### Data availability

Gene expression datasets are available at GSE270570.

To review GEO accession GSE270570:

https://urldefense.com/v3/_https://www.ncbi.nlm.nih.gov/geo/query/acc.cgi?acc=GSE270570;!!MznTZTSvDXGV0Co!BifvZhm5kHoZEkIv4AqUG5m7TYKtQrO4Dg2IYqBPzFPjaAtpCfzGJ01L3ZtuKhfRIwKqPZxfQPjMmyooOUABwYfmFUV7eyxtiDE$

Enter token anadawoqjlgxtod into the box

## ACKNOWLEDGEMENTS

This work was supported by grants from the National Institutes of Health (U54CA231649 to DGM, R35GM124958, R01HD109239 to LAB), Cancer Prevention and Research Institute of Texas (RR140084, RP190414, DGM), UT Southwestern Disease Oriented Scholars Program (DGM), UT Southwestern Endowed Scholars Program (LAB), St. Baldrick’s Foundation Scholar Award (DGM), the 1Million4Anna Foundation (DGM), Welch Foundation (I-2025, LAB), and the American Cancer Society (134230-RSG-20-043-01-DMC, LAB).

## AUTHOR CONTRIBUTIONS

JHM - designed and performed experiments, co-wrote the manuscript

ABE - designed and performed experiments

FV - performed experiments

XB - performed experiments

JK - performed bioinformatics analyses of datasets

YX - supervised bioinformatics analyses performed by JK

PS - performed experiments and analyzed experimental data

RO - performed experiments and bioinformatics analyses

LB - analyzed experimental data, supervised experiments and analyses performed by PS and RO

JK - performed experiments related to xenograft studies

NW - designed and performed animal studies, supervised experiments performed by JK

DGM - conceived project, analyzed experimental data, supervised experiments performed by JHM, ABE, FV and XB, co-wrote the manuscript.

## METHODS

### Cell culture

A673 (CRL-1598), SK-N-MC (HTB-10) were purchased from ATCC and TC32 (Children’s Oncology Group) were cultured in RPMI 1640 media (Sigma-Aldrich R7256) with 10% FBS (), 1% Pen-Strep (Sigma P4332), 1% L-glutamine (Sigma-Aldrich G7513). HEK 293-F cells (Thermo R79007) were cultured in DMEM (Sigma-Aldrich D6429) with 10% FBS, 1% Pen-Strep (Sigma P4332), and 1% L-glutamine (Sigma-Aldrich G7513). All cell lines were maintained in an incubator at 37°C with 5% CO2. Cell lines were periodically tested for mycoplasma and had their identities verified by STR profiling.

### Cell cycle analysis

Cell cycle status was determined using Guava Cell Cycle reagent (Luminex 4500-0220) according to the manufacturer’s instructions. Briefly, cells were washed with PBS, fixed using 70% ethanol, and incubated with cell cycle reagent for 30 min at room temperature. Flow cytometry analysis was performed using Guava easyCyte HT Flow Cytometer and analyzed with Guava InCyte software (Millipore).

### EWSR1-FLI1 degron cell line generation

Degron lines were generated by transfecting cells using the Lipofectamine 3000 (Invitrogen) protocol. Briefly, 1 million cells were plated into each well of a 6-well dish and transfected with 2.5ug of DNA (1:1 repair template plasmid to guide vector) on the same day as plating. 72 hours after transfection, cells were moved from the 6-well to a 10-cm dish, allowed to recover for 24 hours before antibiotic selection (3ug/ml of BSD or 400ug/ml NEO). Clones were isolated and genotyped to ensure correct insertion of repair template.

### Generation of EWSR1-FLI1 expression plasmid and lentivirus production

We constitutively expressed EWSR1-FLI1 constructs using a CMV driven pLVX-IRES-puro (Clontech) backbone. We used EWSR1-FLI1 gene blocks (IDT) as template for expression construct. Plasmid inserts were amplified using Cloneamp HiFi PCR Mix (ClonTech 6329298) or Kapa HiFi HS Mix (KapaBiosystems KK2602). Plasmids were assembled with NEB Assembly Mastermix (NEB E2621X). Plasmids were transformed into Stbl3 E. coli, isolated as single colonies, and sequence verified. Lentivirial plasmids were generated by transfecting 293-F cells with pLVX-EWSR1-FLI1-IRES-Puro plasmid, psPAX2 (Addgene plasmid #12260), and pMD2.G (Addgene plasmid #12259) in a ratio (4:3:1) using TransIT®-LT1Transfection reagent (MIR 2304, Mirus Bio) as described by manufacturer. TC32^EF;SMASh^ cells were transduced with pLVX-EWSR1-FLI1-IRES-Puro lentivirus and selected with puromycin (Sigma P8833).

### Ewing sarcoma cell doubling assay

Cell lines were plated in triplicate for each treatment condition in 6cm plates. Cells were serially passed and counted every 3 days using the ViCell XR cell counter (Beckman Coulter).

### CUT&RUN

CUT&RUN [46] was performed with adjustments made for crosslinking. Briefly, 500,000 nuclei were cryopreserved in Wash150 (20 mM HEPES pH 7.5, 150 mM NaCl, 10 mM NaButyrate, protease inhibitor cocktail (Roche), 0.5 mM Spermidine) + 10% DMSO then stored in liquid nitrogen until experiment. Nuclei were bound to CUTANA Concanavalin A Beads (Epicypher 21-1401) for 15 min, then incubated with 50 μL Wash150 + 0.1% BSA, 2 mM EDTA, and 1 μL primary antibody overnight at 4 °C. Nuclei were resuspended in 100 μL Wash150 + 1 μL secondary antibody at room temperature for 1 h. Nuclei were washed twice in 1 mL Wash150 (with no Spermidine), then resuspended in 200 μL Wash150 (no Spermidine) + 0.2% formaldehyde for 2 min and then quenched with 70.5μL 1M Tris-HCl (pH 8), final concentration of 150mM. Nuclei were washed once in 1 mL Wash350 (20 mM HEPES pH 7.5, 350 mM NaCl, 10 mM NaButyrate, 0.025% Digitonin, protease inhibitor cocktail (Roche), 0.5 mM Spermidine) then incubated in 47.5 μL Wash350 + 2.5 μL pAG-MNase (Epicypher 15-1016) for 1 h. Nuclei were washed twice in 1 mL Wash500 (20 mM HEPES pH 7.5, 500 mM NaCl, 10 mM NaButyrate, 0.025% Digitonin, protease inhibitor cocktail (Roche), 0.5 mM Spermidine), once in 1 mL WashLiCl (20 mM HEPES pH 7.5, 250 mM LiCl, 10 mM NaButyrate, 0.025% Digitonin, protease inhibitor cocktail (Roche), 0.5 mM Spermidine), twice in Wash150 (20 mM HEPES pH 7.5, 150 mM NaCl, 10 mM NaButyrate, 0.025% Digitonin, protease inhibitor cocktail (Roche), 0.5 mM Spermidine) then resuspended in 50 μL Wash150 + 10 mM CaCl2 and incubated for 1 h at 0°C (on aluminum block). Reaction stop and fragment purification is as previously described. Library prep was performed using NEBNext® Ultra™ II for DNA Library Prep using the following protocol (https://www.protocols.io/view/library-prep-for-cut-amp-run-with-nebnext-ultra-ii-kxygxm7pkl8j/v2). The quality of the libraries was assessed using a D1000 ScreenTape on a 2200 TapeStation (Agilent) and quantified using a Qubit dsDNA HS Assay Kit (Thermo Fisher). Libraries with unique adaptor barcodes were multiplexed and sequenced on an Illumina NextSeq 500 (paired-end, 50 base pair reads). Typical sequencing depth was at least 12 million reads per sample.

### CUT&RUN analysis

Raw CUT&RUN reads were adapter and quality trimmed using Trimgalore (47). Trimmed reads were aligned to the human (hg38) reference genome with Bowtie2 (48) (bowtie2 -q -R 3 -N 1 -L 20 -i S,1,0.50 --end-to-end --dovetail --no-mixed -X 2000). Multimapping reads were randomly assigned. Optical duplicate reads were identified and removed using Picard. Reads which mapped to the mitochondrial genome were removed with Samtools (49). Deduplicated bam files were then downsampled according to read depth and merged using Picard. Peak calling was performed with MACS2 software (50) (--keep-dup 10 --nomodel-f BAMPE and an FDR cutoff of 1e-5). Peaks which intersected blacklisted high-signal genomic regions were removed. BigWig files were generated from alignments using deepTools (51) and normalized to counts per million (CPM). Visualization of bigWigs was done in Integrative Genomics Viewer (52). Intersections between different peak sets were made using BEDTools (53, 54). Heatmaps and average profiles were generated using deepTools. Motif enrichment of peak summits was performed using Homer (55).

### RNA preparation

RNA was isolated using Trizol reagent (Thermo Fisher 15596018) and the Directzol RNA Midi Prep Plus Kit (Zymo R2071) according to the manufacturer’s instructions. RNA samples QC, library prep, and sequencing were done by BGI.

### Western blot analysis

Western-blotting was performed using standard methods. Immobilon-P PVDF membranes (Milipore) were used for protein transfer and then blocked using 5% milk in PBS-T (0.1%) for 1h at RT. Primary antibodies were incubated overnight at 4°C in 5% BSA in PBS-Tween (0.1%). Antibodies used were anti-FLI1 rabbit [ERP4646] mAb (Abcam #133485), and anti-β-actin (8H10D10) mouse mAb (#3700, Cell Signaling Technologies). Membranes were washed with PBS-Tween (0.1%) three times for 5 minutes each wash. Secondary antibodies were incubated for 40 min at RT using HRP Linked Horse anti-rabbit IgG (H+L) (CST), and HRP linked anti-mouse IgG (H+L) (CST #7076), at dilution 1:10,000 in 2% milk in PBS-T (0.1%).

### Evaluation of xenograft growth, auxin and danoprevir pharmacokinetics

Animal work described in this manuscript has been approved and conducted under the oversight of the UT Southwestern Institutional Animal Care and Use Committee. UT Southwestern uses the “Guide for the Care and Use of Laboratory Animals” when establishing animal research standards. Xenografts were generated as previously described (Ambati et al 2013). Nod scid mice were injected with 2 million cells @ 1:1 mix with Corning Matrigel Matrix (CLS354234). Xenografts were measured every 3 days until palpable tumors were identified. Tumor-bearing mice were treated with 30 mg/kg danoprevir in vehicle (10% DMSO, 10% PEG400, 80% 0.1M sodium carbonate buffer, pH 10) IP twice daily for 4 days and 3 hours after final dose, mice were euthanized and plasma and tissues collected. In separate studies, additional tumor-bearing mice were treated q12 with 200 mg/kg IAA (3-indole acetic acid, Sigma) formulated in 10% DMSO, 10% Kolliphor EL (Sigma-Aldrich C5135), 80% 0.1M sodium carbonate buffer (pH 9.5). Mice were sacrificed 6, 12 or 24 h after their final dose of either vehicle or IAA. Extracted tumors were divided into pieces for snap freezing in liquid nitrogen or formalin fixation. Frozen tumor pieces were ground into a fine powder using a mortar and pestle and resuspended in RIPA buffer and homogenized for analysis of EWS-FLI1 levels.

Danoprevir levels were monitored by LC-MS/MS using an AB Sciex (Framingham, MA) 3200 QTRAP mass spectrometer coupled to a Shimadzu (Columbia, MD) Prominence LC. Danoprevir was detected with the mass spectrometer in positive ESI MRM (multiple reaction monitoring) mode by following the precursor to fragment ion transition 732.3 to 632.3. An Agilent C18 XDB column (5 micron, 50 x 4.6 mm) was used for chromatography with the following conditions: Buffer A: dH20 + 0.1% formic acid, Buffer B: acetonitrile + 0.1% formic acid with gradient conditions: 0 - 1.0 min 45% B, 1.0 - 4.0 min gradient to 100% B, 4.0 - 5.3 min 100% B, 5.3 - 5.5 min gradient to 45% B, 5.5 - 7.0 min 45% B. Tolbutamide (transition 271.2 to 91.2) from Sigma (St. Louis, MO) was used as an internal standard (IS). At the indicated times post-dose, animals were euthanized and blood collected using acidified citrate dextrose (ACD) anticoagulant and tissues removed, rinsed in PBS, weighed and snap frozen in liquid nitrogen. Liver and tumor tissues were homogenized in a 4x weight by volume of PBS using a BeadBug microtube homogenizer (Millipore Sigma) run for two minutes at 2800 rpm and BeadBug prefilled tubes with 3.0 mm zirconium beads (Sigma Cat #Z763802). Standards were made by spiking blank plasma or tissue homogenate with varying concentrations of danoprevir and processing as for samples. Samples and standards were mixed with a 3-fold volume of acetonitrile containing 0.133% formic acid and 66.7 ng/ml tolbutamide IS, vortexed for 15 seconds, incubated at RT for 10 min and then centrifuged at 16,100 x g for 5 minutes. Supernatant was spun a second time and the resulting supernatant analyzed by LC-MS/MS as described above. Using literature values for the volume of blood in the liver and the measured plasma concentration of danoprevir, liver tissue levels were corrected to remove drug in vasculature^40^.

IAA levels were similarly monitored using an AB Sciex 4000 QTRAP coupled to a Shimadzu Prominence LC. IAA was detected with the mass spectrometer in positive ESI MRM mode by following the precursor to fragment ion transition 173.8 to 127.9. The Agilent C18 XDB column (5 micron, 50 x 4.6 mm) was used for chromatography with the following conditions: Buffer A: dH20 + 0.1% formic acid, 2 mM NH4 acetate; Buffer B: methanol + 0.1% formic acid, 2 mM NH4 acetate with gradient conditions: 0 - 1.0 min 5% B, 1.0 - 2.0 min gradient to 100% B, 2.0 – 3.0 min 100% B, 3.0 – 3.1 min gradient to 5% B, 3.1 – 4.5 min 5% B. After the final indicated dose of IAA, animals were euthanized and blood and tumors harvested as described above. Tumors were homogenized as described above using the BeadBug homogenizer. Plasma samples were diluted 1:10 or 1:100 into 10% mouse plasma in PBS while tumors were diluted 1:5 or 1:10 into 20% blank tumor homogenate in PBS. Standards were prepared in either 10% blank plasma or 20% blank tumor homogenate by spiking these matrices with known amounts of IAA. Diluted samples with mixed 1:1 with methanol containing 0.2% formic acid and 4 mM NH4 acetate containing 100 ng/mL tolbutamide IS, vortexed 15 seconds, incubated at RT for 10 min and centrifuged twice at 16,100 x g. Supernatant was evaluated as described above by LC-MS/MS.

### Bioinformatics

Trim Galore (https://www.bioinformatics.babraham.ac.uk/projects/trim_galore/) was used for quality and adapter trimming. The human reference genome sequence and gene annotation data, hg38, were downloaded from the UCSC Genome Browser and the NCBI RefSeq genome database. The quality of RNA-sequencing libraries was assessed by mapping the reads onto human transcript and ribosomal RNA sequences using the Burrows-Wheeler Aligner (BWA, v0.7.17)^41^). STAR (v2.7.10b)^42^ was used to align the reads to the human genome, and SAMtools (v1.16.1)^43^ was used to sort the alignments. The HTSeq Python package^44^ was used to count reads per gene. The DESeq2 R Bioconductor package^45,46^ was employed to normalize read counts and identify differentially expressed (DE) genes. The gene set data for chemical and genetic perturbations (CGP) was downloaded from the Molecular Signatures Database (MSigDB, https://www.gsea-msigdb.org/gsea/msigdb/), and enriched and over-represented gene sets were identified using GSEA software (v4.3.3) ^32^and clusterProfiler ^47^, respectively. *Trim Galore* was developed at The Babraham Institute by @FelixKrueger, now part of [Altos Labs](https://www.altoslabs.com/).

## REFERENCES

1. Gröbner, S.N., Worst, B.C., Weischenfeldt, J., Buchhalter, I., Kleinheinz, K., Rudneva, V.A., Johann, P.D., Balasubramanian, G.P., Segura-Wang, M., Brabetz, S., et al. (2018). The landscape of genomic alterations across childhood cancers. Nature 555, 321–327. 10.1038/nature25480.

2. Ma, X., Liu, Y., Liu, Y., Alexandrov, L.B., Edmonson, M.N., Gawad, C., Zhou, X., Li, Y., Rusch, M.C., Easton, J., et al. (2018). Pan-cancer genome and transcriptome analyses of 1,6*9*9 paediatric leukaemias and solid tumours. Nature 555, 371–376. 10.1038/nature25795.

3. Druker, B.J., Sawyers, C.L., Kantarjian, H., Resta, D.J., Reese, S.F., Ford, J.M., Capdeville, R., and Talpaz, M. (2001). Activity of a Specific Inhibitor of the BCR-ABL Tyrosine Kinase in the Blast Crisis of Chronic Myeloid Leukemia and Acute Lymphoblastic Leukemia with the Philadelphia Chromosome. N. Engl. J. Med. 344, 1038–1042. 10.1056/nejm200104053441402.

4. Druker, B.J., Talpaz, M., Resta, D.J., Peng, B., Buchdunger, E., Ford, J.M., Lydon, N.B., Kantarjian, H., Capdeville, R., Ohno-Jones, S., et al. (2001). Efficacy and Safety of a Specific Inhibitor of the BCR-ABL Tyrosine Kinase in Chronic Myeloid Leukemia. N. Engl. J. Med. 344, 1031–1037. 10.1056/nejm200104053441401.

5. Hadoux, J., Elisei, R., Brose, M.S., Hoff, A.O., Robinson, B.G., Gao, M., Jarzab, B., Isaev, P., Kopeckova, K., Wadsley, J., et al. (2023). Phase 3 Trial of Selpercatinib in Advanced RET-Mutant Medullary Thyroid Cancer. N. Engl. J. Med. 389, 1851–1861. 10.1056/nejmoa2309719.

6. Rabbitts, T.H. (1994). Chromosomal translocations in human cancer. Nature 372, 143–149. 10.1038/372143a0.

7. Delattre, O., Zucman, J., Plougastel, B., Desmaze, C., Melot, T., Peter, M., Kovar, H., Joubert, I., Jong, P. de, Rouleau, G., et al. (1992). Gene fusion with an ETS DNA-binding domain caused by chromosome translocation in human tumours. Nature 359, 162–165. 10.1038/359162a0.

8. Tirode, F., Consortium, for the St.J.C.R.H.U.P.C.G.P. and the I.C.G., Surdez, D., Ma, X., Parker, M., Deley, M.C.L., Bahrami, A., Zhang, Z., Lapouble, E., Grossetête-Lalami, S., et al. (2014). Genomic Landscape of Ewing Sarcoma Defines an Aggressive Subtype with Co-Association of STAG2 and TP53 Mutations. Cancer Discov. 4, 1342–1353. 10.1158/2159-8290.cd-14-0622.

9. Crompton, B.D., Stewart, C., Taylor-Weiner, A., Alexe, G., Kurek, K.C., Calicchio, M.L., Kiezun, A., Carter, S.L., Shukla, S.A., Mehta, S.S., et al. (2014). The Genomic Landscape of Pediatric Ewing Sarcoma. Cancer Discov. 4, 1326–1341. 10.1158/2159-8290.cd-13-1037.

10. Brohl, A.S., Solomon, D.A., Chang, W., Wang, J., Song, Y., Sindiri, S., Patidar, R., Hurd, L., Chen, L., Shern, J.F., et al. (2014). The Genomic Landscape of the Ewing Sarcoma Family of Tumors Reveals Recurrent STAG2 Mutation. PLoS Genet. 10, e1004475. 10.1371/journal.pgen.1004475.

11. Povedano, J.M., Li, V., Lake, K.E., Bai, X., Rallabandi, R., Kim, J., Xie, Y., Brabander, J.K.D., and McFadden, D.G. (2022). TK216 targets microtubules in Ewing sarcoma cells. Cell Chem. Biol. 29, 1325–1332.e4. 10.1016/j.chembiol.2022.06.002.

12. Johnson, K.M., Mahler, N.R., Saund, R.S., Theisen, E.R., Taslim, C., Callender, N.W., Crow, J.C., Miller, K.R., and Lessnick, S.L. (2017). Role for the EWS domain of EWS/FLI in binding GGAA-microsatellites required for Ewing sarcoma anchorage independent growth. Proc. Natl. Acad. Sci. 114, 9870–9875. 10.1073/pnas.1701872114.

13. Boulay, G., Volorio, A., Iyer, S., Broye, L.C., Stamenkovic, I., Riggi, N., and Rivera, M.N. (2018). Epigenome editing of microsatellite repeats defines tumor-specific enhancer functions and dependencies. Genes Dev. 32, 1008–1019. 10.1101/gad.315192.118.

14. Riggi, N., Knoechel, B., Gillespie, S.M., Rheinbay, E., Boulay, G., Suvà, M.L., Rossetti, N.E., Boonseng, W.E., Oksuz, O., Cook, E.B., et al. (2014). EWS-FLI1 Utilizes Divergent Chromatin Remodeling Mechanisms to Directly Activate or Repress Enhancer Elements in Ewing Sarcoma. Cancer Cell 26, 668–681. 10.1016/j.ccell.2014.10.004.

15. Gangwal, K., Sankar, S., Hollenhorst, P.C., Kinsey, M., Haroldsen, S.C., Shah, A.A., Boucher, K.M., Watkins, W.S., Jorde, L.B., Graves, B.J., et al. (2008). Microsatellites as EWS/FLI response elements in Ewing’s sarcoma. Proc. Natl. Acad. Sci. 105, 10149–10154. 10.1073/pnas.0801073105.

16. Guillon, N., Tirode, F., Boeva, V., Zynovyev, A., Barillot, E., and Delattre, O. (2009). The Oncogenic EWS-FLI1 Protein Binds In Vivo GGAA Microsatellite Sequences with Potential Transcriptional Activation Function. PLoS ONE 4, e4932. 10.1371/journal.pone.0004932.

17. Prieur, A., Tirode, F., Cohen, P., and Delattre, O. (2004). EWS/FLI-1 Silencing and Gene Profiling of Ewing Cells Reveal Downstream Oncogenic Pathways and a Crucial Role for Repression of Insulin-Like Growth Factor Binding Protein 3. Mol. Cell. Biol. 24, 7275–7283. 10.1128/mcb.24.16.7275-7283.2004.

18. Hu, H., Zielinska-Kwiatkowska, A., Munro, K., Wilcox, J., Wu, D.Y., Yang, L., and Chansky, H.A. (2008). EWS/FLI1 suppresses retinoblastoma protein function and senescence in Ewing’s sarcoma cells. J. Orthop. Res. 26, 886–893. 10.1002/jor.20597.

19. Smith, R., Owen, L.A., Trem, D.J., Wong, J.S., Whangbo, J.S., Golub, T.R., and Lessnick, S.L. (2006). Expression profiling of EWS/FLI identifies NKX2.2 as a critical target gene in Ewing’s sarcoma. Cancer Cell 9, 405–416. 10.1016/j.ccr.2006.04.004.

20. Mateo-Lozano, S., Gokhale, P.C., Soldatenkov, V.A., Dritschilo, A., Tirado, O.M., and Notario, V. (2006). Combined Transcriptional and Translational Targeting of EWS/FLI-1 in Ewing’s Sarcoma. Clin. Cancer Res. 12, 6781–6790. 10.1158/1078-0432.ccr-06-0609.

21. Chansky, H.A., Barahmand-pour, F., Mei, Q., Kahn-Farooqi, W., Zielinska-Kwiatkowska, A., Blackburn, M., Chansky, K., Conrad, E.U., Bruckner, J.D., Greenlee, T.K., et al. (2004). Targeting of EWS/FLI-1 by RNA interference attenuates the tumor phenotype of Ewing’s sarcoma cells in vitro. J. Orthop. Res. 22, 910–917. 10.1016/j.orthres.2003.12.008.

22. Matsunobu, T., Tanaka, K., Nakamura, T., Nakatani, F., Sakimura, R., Hanada, M., Li, X., Okada, T., Oda, Y., Tsuneyoshi, M., et al. (2006). The Possible Role of EWS-Fli1 in Evasion of Senescence in Ewing Family Tumors. Cancer Res. 66, 803–811. 10.1158/0008-5472.can-05-1972.

23. Nakatani, F., Tanaka, K., Sakimura, R., Matsumoto, Y., Matsunobu, T., Li, X., Hanada, M., Okada, T., and Iwamoto, Y. (2003). Identification of p21 WAF1/CIP1 as a Direct Target of EWS-Fli1 Oncogenic Fusion Protein*. J. Biol. Chem. 278, 15105–15115. 10.1074/jbc.m211470200.

24. Martinez-Lage, M., Torres-Ruiz, R., Puig-Serra, P., Moreno-Gaona, P., Martin, M.C., Moya, F.J., Quintana-Bustamante, O., Garcia-Silva, S., Carcaboso, A.M., Petazzi, P., et al. (2020). In vivo CRISPR/Cas9 targeting of fusion oncogenes for selective elimination of cancer cells. Nat. Commun. 11, 5060. 10.1038/s41467-020-18875-x.

25. Nabet, B., Ferguson, F.M., Seong, B.K.A., Kuljanin, M., Leggett, A.L., Mohardt, M.L., Robichaud, A., Conway, A.S., Buckley, D.L., Mancias, J.D., et al. (2020). Rapid and direct control of target protein levels with VHL-recruiting dTAG molecules. Nat. Commun. 11, 4687. 10.1038/s41467-020-18377-w.

26. Saito, Y., and Kanemaki, M.T. (2021). Targeted Protein Depletion Using the Auxin-Inducible Degron 2 (AID2) System. Curr. Protoc. 1, e219. 10.1002/cpz1.219.

27. Yesbolatova, A., Saito, Y., Kitamoto, N., Makino-Itou, H., Ajima, R., Nakano, R., Nakaoka, H., Fukui, K., Gamo, K., Tominari, Y., et al. (2020). The auxin-inducible degron 2 technology provides sharp degradation control in yeast, mammalian cells, and mice. Nat. Commun. 11, 5701. 10.1038/s41467-020-19532-z.

28. Chung, H.K., Jacobs, C.L., Huo, Y., Yang, J., Krumm, S.A., Plemper, R.K., Tsien, R.Y., and Lin, M.Z. (2015). Tunable and reversible drug control of protein production via a self-excising degron. Nat. Chem. Biol. 11, 713–720. 10.1038/nchembio.1869.

29. Arvand, A., Welford, S.M., Teitell, M.A., and Denny, C.T. (2001). The COOH-terminal domain of FLI-1 is necessary for full tumorigenesis and transcriptional modulation by EWS/FLI-1. Cancer Res. 61, 5311–5317.

30. Boone, M.A., Taslim, C., Crow, J.C., Selich-Anderson, J., Byrum, A.K., Showpnil, I.A., Sunkel, B.D., Wang, M., Stanton, B.Z., Theisen, E.R., et al. (2021). The FLI portion of EWS/FLI contributes a transcriptional regulatory function that is distinct and separable from its DNA-binding function in Ewing sarcoma. Oncogene 40, 4759–4769. 10.1038/s41388-021-01876-5.

31. Gierisch, M.E., Pfistner, F., Lopez-Garcia, L.A., Harder, L., Schäfer, B.W., and Niggli, F.K. (2016). Proteasomal Degradation of the EWS-FLI1 Fusion Protein Is Regulated by a Single Lysine Residue*. J. Biol. Chem. 291, 26922–26933. 10.1074/jbc.m116.752063.

32. Subramanian, A., Tamayo, P., Mootha, V.K., Mukherjee, S., Ebert, B.L., Gillette, M.A., Paulovich, A., Pomeroy, S.L., Golub, T.R., Lander, E.S., et al. (2005). Gene set enrichment analysis: A knowledge-based approach for interpreting genome-wide expression profiles. Proc. Natl. Acad. Sci. 102, 15545–15550. 10.1073/pnas.0506580102.

33. Mootha, V.K., Lindgren, C.M., Eriksson, K.-F., Subramanian, A., Sihag, S., Lehar, J., Puigserver, P., Carlsson, E., Ridderstråle, M., Laurila, E., et al. (2003). PGC-1α-responsive genes involved in oxidative phosphorylation are coordinately downregulated in human diabetes. Nat. Genet. 34, 267–273. 10.1038/ng1180.

34. Gao, Y., He, X.-Y., Wu, X.S., Huang, Y.-H., Toneyan, S., Ha, T., Ipsaro, J.J., Koo, P.K., Joshua-Tor, L., Bailey, K.M., et al. (2023). ETV6 dependency in Ewing sarcoma by antagonism of EWS-FLI1-mediated enhancer activation. Nat. Cell Biol. 25, 298–308. 10.1038/s41556-022-01060-1.

35. Lu, D.Y., Ellegast, J.M., Ross, K.N., Malone, C.F., Lin, S., Mabe, N.W., Dharia, N.V., Meyer, A., Conway, A., Su, A.H., et al. (2023). The ETS transcription factor ETV6 constrains the transcriptional activity of EWS–FLI to promote Ewing sarcoma. Nat. Cell Biol. 25, 285–297. 10.1038/s41556-022-01059-8.

36. Seiwert, S.D., Andrews, S.W., Jiang, Y., Serebryany, V., Tan, H., Kossen, K., Rajagopalan, P.T.R., Misialek, S., Stevens, S.K., Stoycheva, A., et al. (2008). Preclinical Characteristics of the Hepatitis C Virus NS3/4A Protease Inhibitor ITMN-191 (R7227). Antimicrob. Agents Chemother. 52, 4432–4441. 10.1128/aac.00699-08.

37. Chaturvedi, A., Hoffman, L.M., Welm, A.L., Lessnick, S.L., and Beckerle, M.C. (2012). The EWS/FLI Oncogene Drives Changes in Cellular Morphology, Adhesion, and Migration in Ewing Sarcoma. Genes Cancer 3, 102–116. 10.1177/1947601912457024.

38. Theisen, E.R., Miller, K.R., Showpnil, I.A., Taslim, C., Pishas, K.I., and Lessnick, S.L. (2019). Transcriptomic analysis functionally maps the intrinsically disordered domain of EWS/FLI and reveals novel transcriptional dependencies for oncogenesis. Genes Cancer 10, 21–38. 10.18632/genesandcancer.188.

39. Boulay, G., Sandoval, G.J., Riggi, N., Iyer, S., Buisson, R., Naigles, B., Awad, M.E., Rengarajan, S., Volorio, A., McBride, M.J., et al. (2017). Cancer-Specific Retargeting of BAF Complexes by a Prion-like Domain. Cell 171, 163–178.e19. 10.1016/j.cell.2017.07.036.

40. Kwon, Y. (2002). Handbook of Essential Pharmacokinetics, Pharmacodynamics and Drug Metabolism for Industrial Scientists. 10.1007/b112416.

41. Li, H., and Durbin, R. (2009). Fast and accurate short read alignment with Burrows–Wheeler transform. Bioinformatics 25, 1754–1760. 10.1093/bioinformatics/btp324.

42. Dobin, A., Davis, C.A., Schlesinger, F., Drenkow, J., Zaleski, C., Jha, S., Batut, P., Chaisson, M., and Gingeras, T.R. (2012). STAR: ultrafast universal RNA-seq aligner. Bioinformatics 29, 15–21. 10.1093/bioinformatics/bts635.

43. Li, H., Handsaker, B., Wysoker, A., Fennell, T., Ruan, J., Homer, N., Marth, G., Abecasis, G., Durbin, R., and Subgroup, 1000 Genome Project Data Processing (2009). The Sequence Alignment/Map format and SAMtools. Bioinformatics 25, 2078–2079. 10.1093/bioinformatics/btp352.

44. Anders, S., Pyl, P.T., and Huber, W. (2014). HTSeq—a Python framework to work with high-throughput sequencing data. Bioinformatics 31, 166–169. 10.1093/bioinformatics/btu638.

45. Gentleman, R.C., Carey, V.J., Bates, D.M., Bolstad, B., Dettling, M., Dudoit, S., Ellis, B., Gautier, L., Ge, Y., Gentry, J., et al. (2004). Bioconductor: open software development for computational biology and bioinformatics. Genome Biol. 5, R80. 10.1186/gb-2004-5-10-r80.

46. Anders, S., and Huber, W. (2010). Differential expression analysis for sequence count data. Genome Biol. 11, R106. 10.1186/gb-2010-11-10-r106.

47. Yu, G., Wang, L.-G., Han, Y., and He, Q.-Y. (2012). clusterProfiler: an R Package for Comparing Biological Themes Among Gene Clusters. OMICS: A J. Integr. Biol. 16, 284–287. 10.1089/omi.2011.0118.

