## Supplemental Figures for "Endogenous EWSR1-FLI1 degron alleles enable control of fusion oncoprotein expression in tumor cell lines and xenografts"

#### Figure S1. Degron tags enable depletion of endogenous EWSR1-FLI1

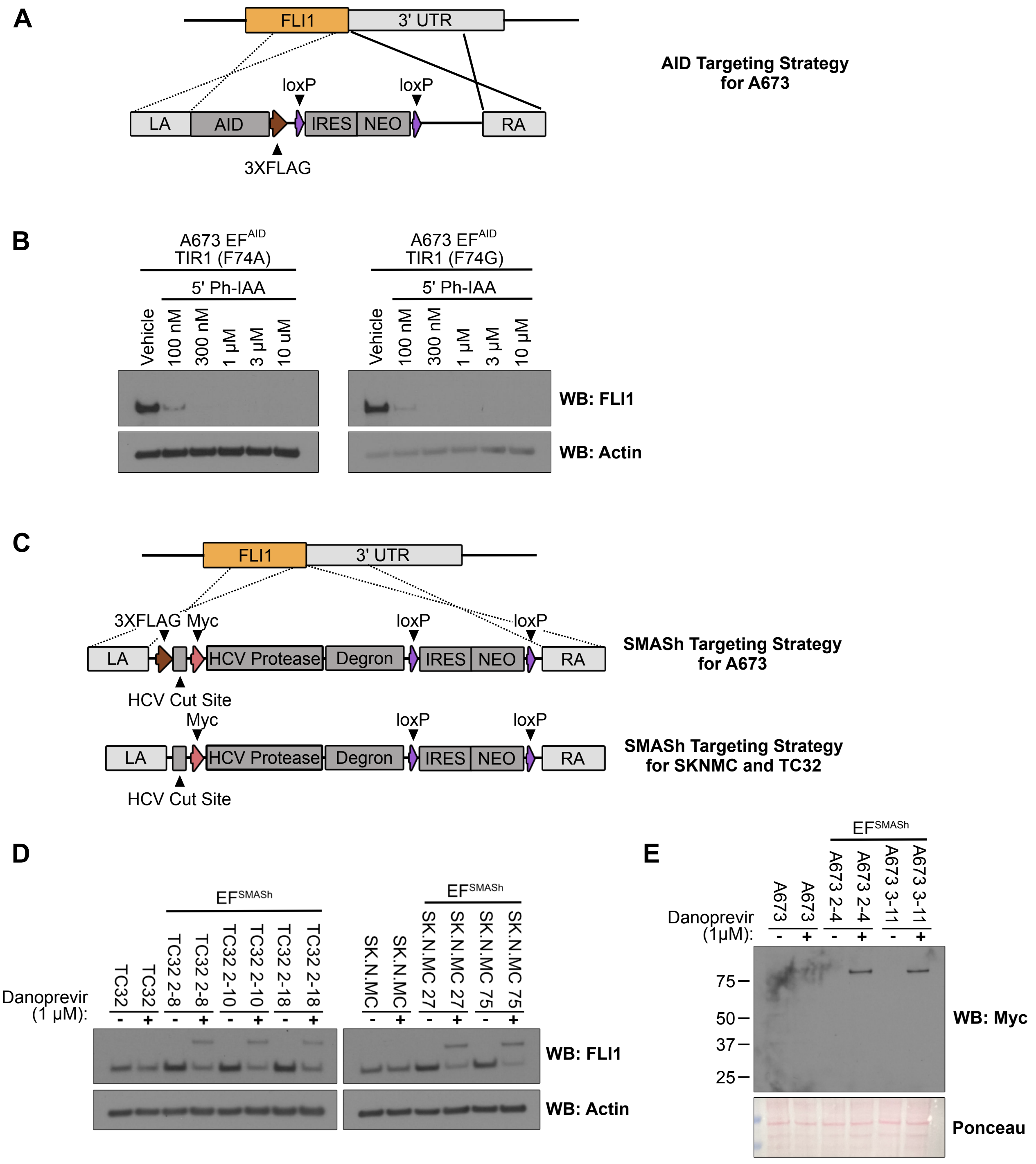


**Figure S1: Depletion of endogenous EWSR1-FLI1 with inducible degron systems.** A) Schematic depicting CRISPR/Cas9 mediated knock-in of AID cassette into the EWS-FLI1 locus. B) Immunoblot for EWSR1-FLI1 (FLI1) in indicated cell lines. Cell lines were exposed to either DMSO or 5-Ph-IAA at the indicated concentration for 24 hours prior to collection. C) Schematic depicting CRISPR/Cas9 mediated knock-in of SMASh cassette into the EWSR1-FLI1 locus. D) Immunoblot for EWSR1-FLI1 (FLI1) in indicated cell lines. Cell lines were exposed to either DMSO or Danoprevir (1 µM) for 24 hours prior to collection. D) Immunoblot for Myc epitope tag in indicated cell lines. Cell lines were exposed to either DMSO or Danoprevir (1 µM) for 24 hours prior to collection.

#### Figure S2. C-terminal AID tag on EWSR1-FLI1 does not disrupt DNA binding

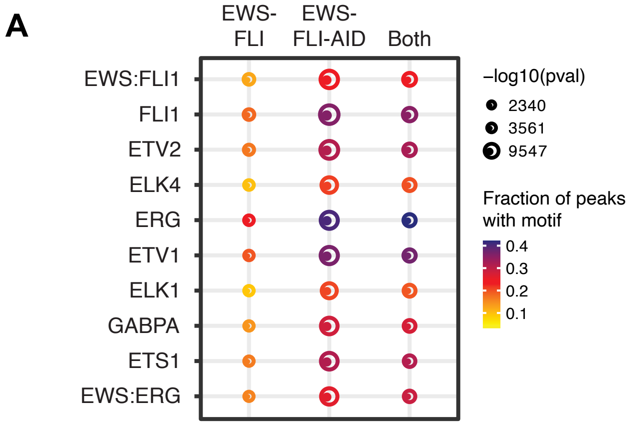


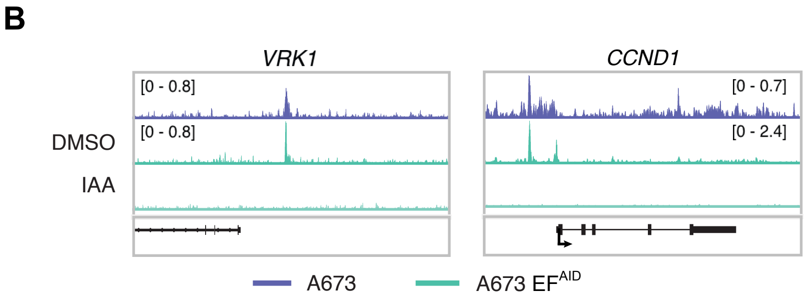


**Figure S2: C-terminal AID tag on EWSR1-FLI1 does not alter DNA binding** A) Motif analysis of transcription factor binding sites identified in wild-type A673 cells, A673 $EF^{AID}$ cells, or in both datasets. Note that EWS:FLI binding site represents GGAA repeat microsatellite sequences. B) Genome browser representations of FLI CUT&RUN in A673 cells and A673 $EF^{AID}$ cells treated with either DMSO or IAA (100μM) for 24 hrs. The y-axis represents read density in reads per million mapped reads (rpm).

### Figure S3. EWSR1-FLI1 depletion induces G1/S arrest

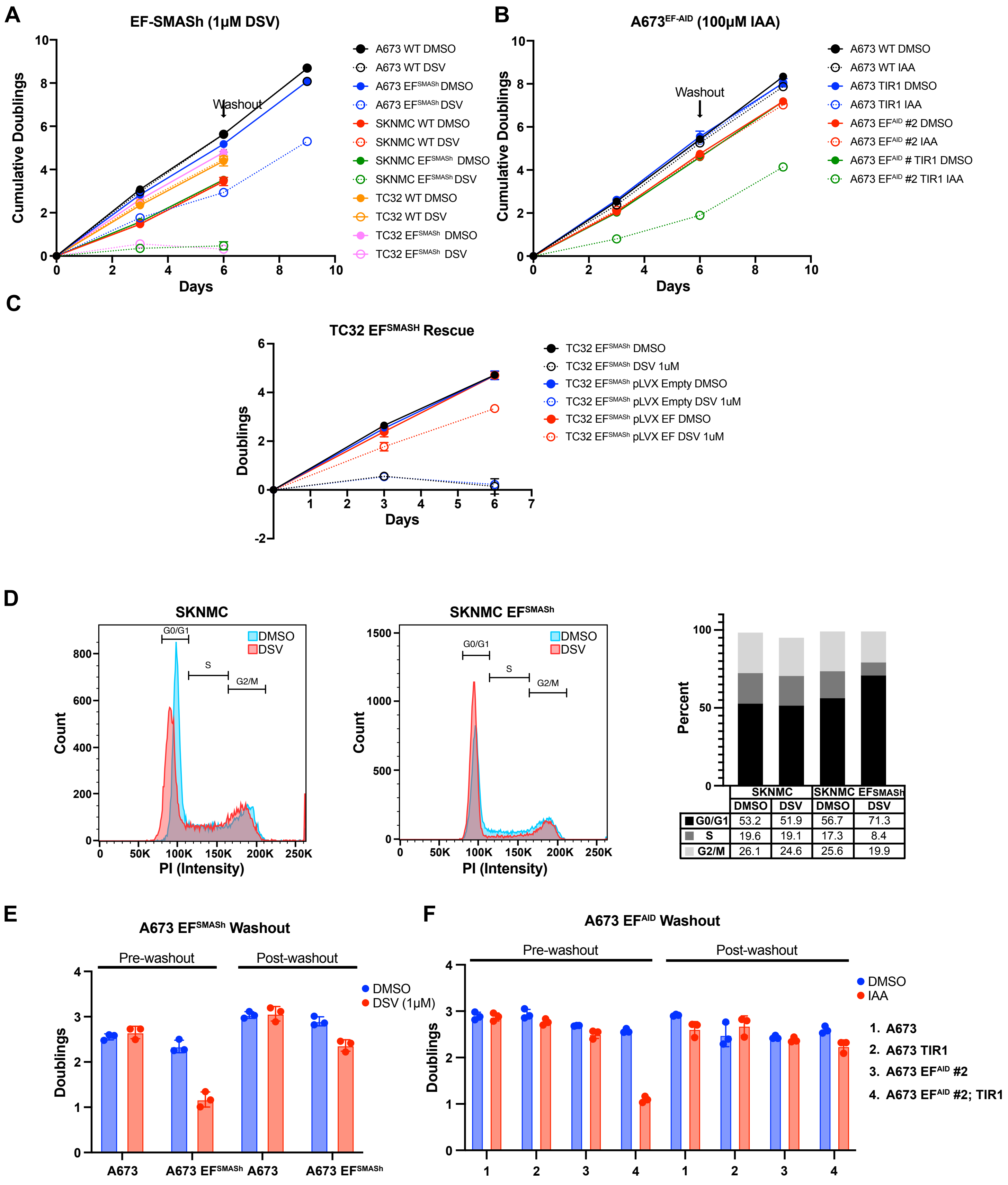


**Figure S3: EWSR1-FLI1 depletion leads to G1/S arrest** A-C) Growth curves for indicated cell lines treated with either vehicle, 1 μM Danoprevir or IAA (summary of data at day 6 for each plot is shown in Figure 3A-B and 3D). D) Cell cycle analysis using propidium iodine stain. Flow cytometry plots (left two panels) for indicated cell lines treated with either vehicle or 1 μM Danoprevir for 72 hours before cells were collected. Plot (left panel) of cells (percentage) in each phase of the cell cycle based on flow cytometry data. For E) and F) Data from panel A and B, respectively, is summarized as cumulative doublings. Pre-washout refers to doublings between day 3 and 6, post-washout refers to doublings between day 6 and 9.

#### Figure S4. Core set of EWSR1-FLI1 response genes shared across EWS cell lines

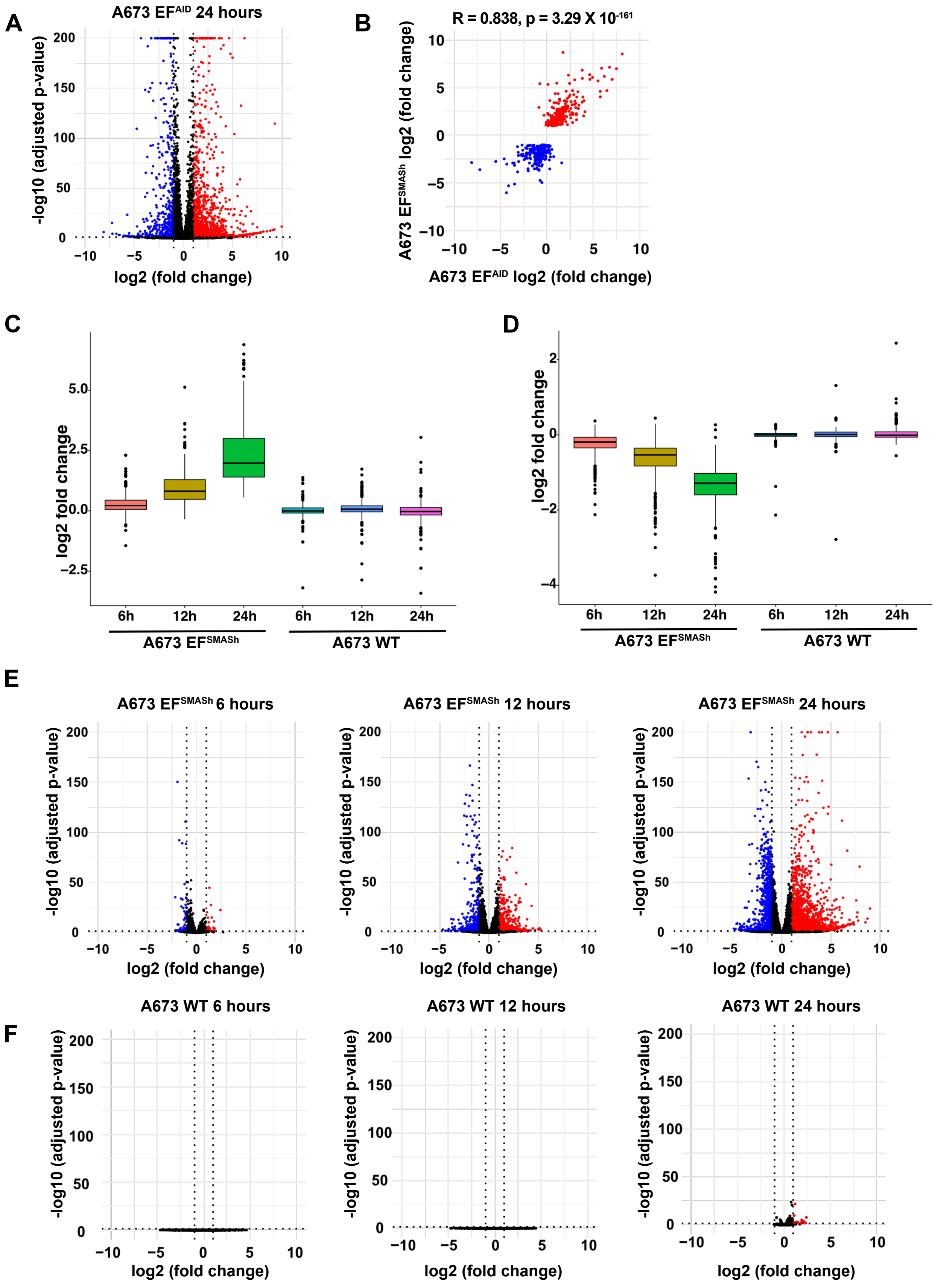


**Figure S4: Core set of EWSR1-FLI1 response genes shared across Ewing sarcoma cell lines** A) Volcano plots for RNA sequencing of A673 $EF^{AID}$ cells following treatment with IAA for 24 hours. Significantly induced transcripts labeled in red, and significantly down-regulated transcripts labeled in blue .B) Plot comparing differentially expressed genes from A673 $EF^{SMASh}$ cells and A673 $EF^{AID}$ A673 $EF^{SMASh}$ and A673 $EF^{AID}$ cells treated with danoprevir (1μM) or IAA (100μM) respectively. C-D) Volcano plots from RNA sequencing datasets obtained at the indicated times following treatment of A673 $EF^{SMASh}$ (C) or A673 wild-type (D) cells with 312nM danoprevir.

#### Figure S5. EWSR1-FLI1 is required for tumor maintenance in vivo

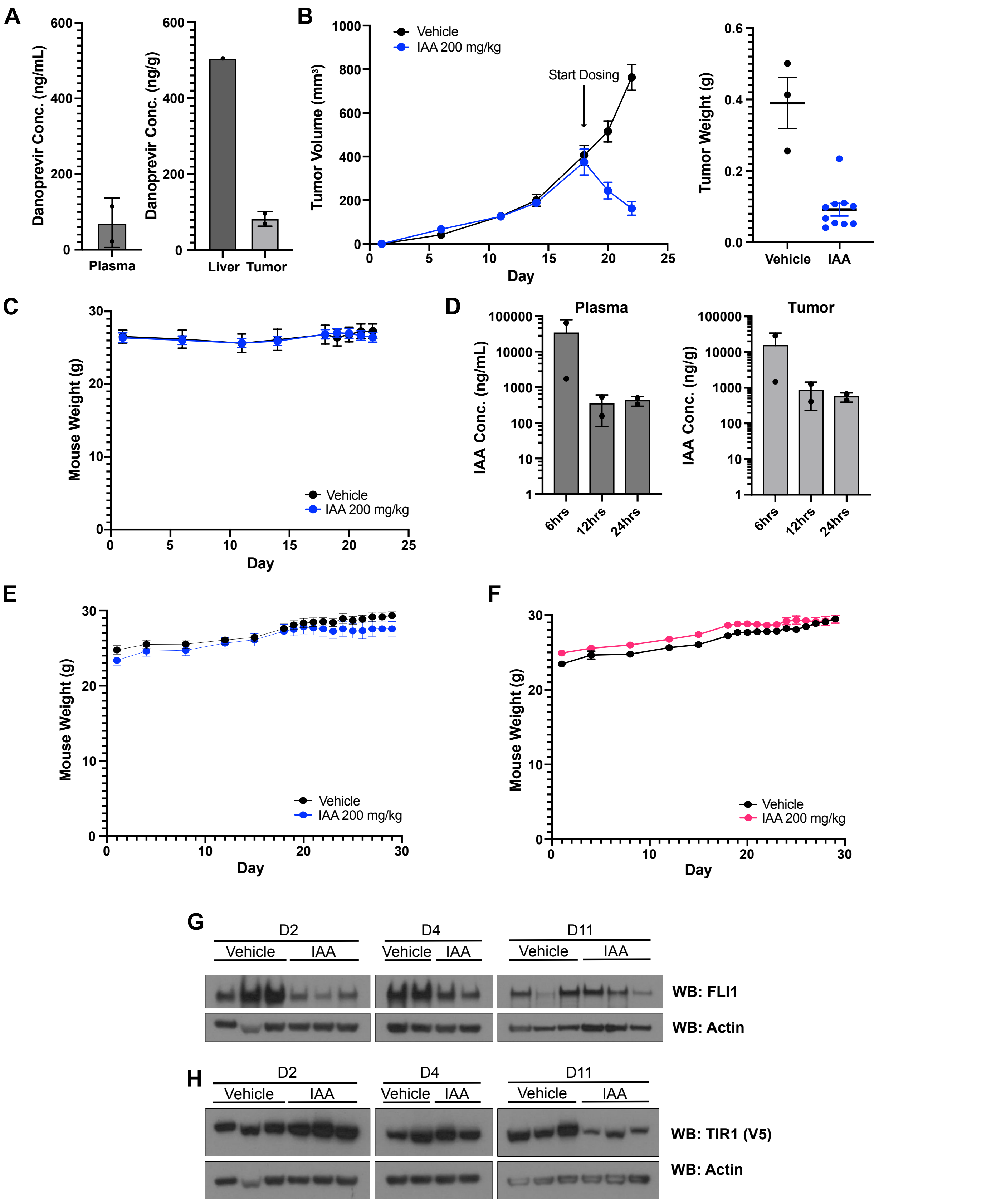


**Figure S5: EWSR1-FLI1 is required for tumor maintenance in vivo** A) danoprevir levels were determined using LC-MS/MS in the plasma, liver and tumor 3 hours after the final dose was administered (n = 2). B) Tumor volume (left) and tumor mass (right) for A673$^{EF;AID:TIR1}$ xenografts treated with vehicle (n = 3) or 200 mg/kg IAA (n = 10). Vehicle or IAA was administered twice daily via IP for a total of 4 days. C) Mouse weight for A673$^{EF;AID:TIR1}$ xenografts (from panel B). D) IAA levels were determined using LC-MS/MS in the plasma and tumor 3, 12 and 24 hours after the final dose of IAA was administered (n = 2 for each time point). E-F) Mouse weight for A673$^{EF;AID:TIR1}$ and A673$^{EF;AID}$ xenografts (from Figure 5A-B). G-H) Immunoblot for EWS-FLI1 (FLI1) or TIR1 (V5) from animals sacrificed 2, 4, or 11 days (D2, D4 and D11 respectively) after IAA treatment was initiated (from Figure 5A).
